# Discovering single-cell eQTLs from scRNA-seq data only

**DOI:** 10.1101/2021.06.10.447906

**Authors:** Tianxing Ma, Haochen Li, Xuegong Zhang

## Abstract

eQTL studies are essential for understanding genomic regulation. Effects of genetic variations on gene regulation are cell-type-specific and cellular-context-related, so studying eQTLs at a single-cell level is crucial. The ideal solution is to use both mutation and expression data from the same cells. However, current technology of such paired data in single cells is still immature. We present a new method, eQTLsingle, to discover eQTLs only with single cell RNA-seq (scRNA-seq) data, without genomic data. It detects mutations from scRNA-seq data and models gene expression of different genotypes with the zero-inflated negative binomial (ZINB) model to find associations between genotypes and phenotypes at single-cell level. On a glioblastoma and gliomasphere scRNA-seq dataset, eQTLsingle discovered hundreds of cell-type-specific tumor-related eQTLs, most of which cannot be found in bulk eQTL studies. Detailed analyses on examples of the discovered eQTLs revealed important underlying regulatory mechanisms. eQTLsingle is a unique powerful tool for utilizing the huge scRNA-seq resources for single-cell eQTL studies, and it is available for free academic use at https://github.com/horsedayday/eQTLsingle.

## 1. Introduction

Genome variations have significant influences on phenotypes[1,2]. Most variations are at single bases, called single-nucleotide variants (SNVs). Some variations have been found responsible for traits such as curly hair[3], tallness[4], and common diseases like diabetes[5], and Alzheimer’s disease[6], etc. A genomic locus associated with variations of downstream gene expression is called an expression quantitative trait locus or eQTL. eQTL studies are important for decoding genomic regulation mechanisms and for improving understandings on disease pathogenesis.

Most existing eQTL studies are based on bulk-level genomic and transcriptomic data, which can only reflect average effects in mixtures of cells. Researchers found that many eQTL effects are cell-type-specific and cellular-context-related[7–9]. Bulk-level analyses cannot detect such eQTLs. We should conduct eQTL analyses at single-cell level. Single-cell sequencing technologies provided great potentials for exploring eQTL effects at single-cell level. Scientists have conducted several eQTL studies using single-cell data recently. For example, Wijst et. al. found multiple personalized cell-type-specific eQTLs by quantifying gene expression levels of peripheral blood mononuclear cells with scRNA-seq data and using corresponding bulk genomic data for detecting mutations[10]. Orozco et. al. associated donors’ genotypes with single-nucleus RNA-seq data, and discovered new putative causal genes for age-related macular degeneration[11]. Cuomo et. al. genotyped hundreds of bulk samples and quantified gene expression with scRNA-seq to study differentiation in early human development[12]. These single-cell studies brought new eQTL discoveries, but the genomic variation data are still at bulk level. They cannot find genomic variations of single cells. For example, mutations in tumor cells are heterogeneous and profoundly influence cellular functions, tumor evolution[13], and treatment responses[14]. Genomic regulations specific to cells or cell-types cannot be revealed with bulk-level genomic data. It’s necessary in many scenarios to do eQTL studies with mutation and expression data both of the single-cell level.

The ideal solution is to sequence the genome and transcriptome in parallel in the same single cells. We have developed a method for eQTL analysis with such paired single-cell sequencing data[15]. However, paired sequencing technologies are still immature, and the genome coverage of current single-cell pair-sequencing data is too shallow for effective eQTL analysis. Several previous works have shown that mutations in gene regions can be detected from RNA-seq data reliably[16,17]. Inspired by this, we propose to use mutations detected from scRNA-seq data to explore their effects on genes. In this paper, we propose a new method eQTLsingle to study eQTLs at a single-cell level using single-cell or single-nucleus RNA-seq data only. It detects mutations in coding regions from scRNA-seq data and therefore gets the mutation and expression profiles in same single cells simultaneously. We apply the ZINB model to fit the gene expression data[18] and associate mutations with the gene expression. The model can depict excess zeros and consider the common drop-out problem in scRNA-seq data well[18]. We applied eQTLsingle on a scRNA-seq dataset of glioblastoma and gliomasphere cells, and discovered hundreds of cell-type-specific cis-eQTLs and trans-eQTLs on genes related with brain tumor. The majority of them (83.57% of the cis-eQTLs and all the trans-eQTLs) have not been reported in the bulk-level GTEx database. Detailed investigations into examples of these results showed the high potential of important discoveries of this single-cell eQTL strategy. eQTLsingle is a unique tool for utilizing the rich single cell RNA-seq data to discover single-cell eQTLs that are crucial for understanding gene regulations behind many cell-type-specific biological processes.

## 2. Result

### 2.1 Overview of eQTLsingle

The ideal solution for single-cell eQTL analysis is to sequence genome and transcriptome for each single-cell simultaneously. Available paired single-cell sequencing technologies, such as G&T-seq[19] and DRseq[20], are still under development. We analyzed some published G&T-seq data and found that coverage of G&T-seq data is very low. Most loci have at most one read in a cell in the data (Supplementary Figure S1). Such shallow coverage is far from sufficient for reliable SNV calling.

We therefore developed the method eQTLsingle to do single-cell eQTL analysis with single-cell or single-nucleus RNA-seq data only. It detects SNVs in exon regions from the same scRNA-seq data as used for calculating gene expression. This gives high-quality genomic variation and gene expression information in the same cells. When we use it to detect eQTLs at single-cell level for specific cell types or cell states, we can divide the cells into the corresponding subsets and conduct eQTL analysis in each subset separately. For an SNV in a subset, we model gene expressions in the mutated cell population (cells with the alternative allele, denoted as the ALT group below) and non-mutated cell population (cells with the reference allele, denoted as the REF group below) with zero-inflated negative binomial (ZINB) models. The ZINB model has been shown effective in fitting scRNA-seq data with drop-out problems[18]. eQTLsingle detects eQTLs, i.e., associations of mutations with gene expression variations, by comparing the ZINB parameters between the ALT and REF groups using likelihood ratio test (LRT). The entire method is illustrated in Figure 1. scRNA-seq data have reads only in the exon regions of the expressed genes. Therefore, eQTLsingle can only detect eQTLs in those regions. But the number of SNVs that can be covered in scRNA-seq data is much larger than the number that can be detected from paired single-cell sequencing data.

**Figure 1.**
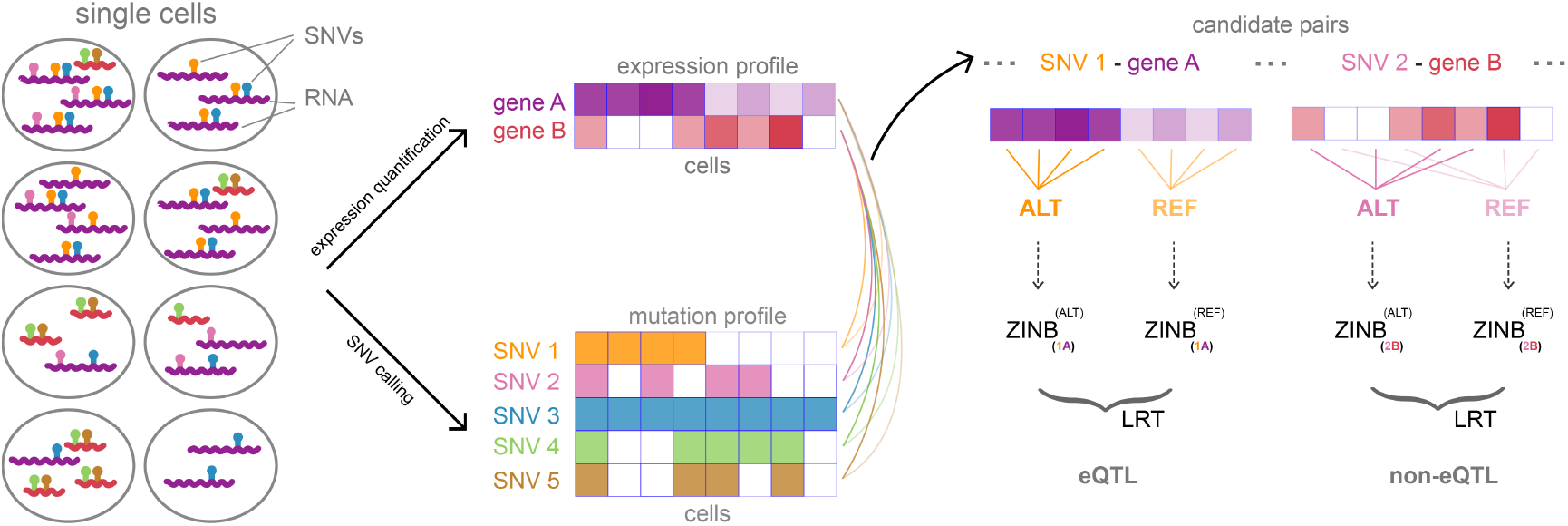
Illustration of the eQTLsingle method. It first calculates gene expression and calls SNVs simultaneously from the raw reads of scRNA-seq data. For each detected SNV, it looks for possible eQTL effect on all candidate target genes by dividing the cells into the REF group or ALT group according to the genotype of cells at this locus first. It then builds a ZINB model for the gene expression in each group, and uses likelihood ratio test (LRT) to compare the models of the two groups of the same gene. The SNV is identified as an eQTL for the target gene if the test finds a significant difference between the two models.

There are two noticeable differences in analyzing SNVs detected from bulk RNA-seq data and scRNA-seq data. Firstly, in scRNA-seq data, loci without mutation reported by a SNV-calling program may be truly with no mutation but also may be due to low read coverage at the loci. For loci without enough reads, we mark them as missing data and do not use them in the eQTL analysis. Secondly, because of possible unbalanced expression of paternal alleles and maternal alleles of genes (the allelic expression) in single cells[21], we cannot distinguish between homogenous and heterogeneous mutations from scRNA-seq data only. So, the genotypes are only categorized as ALT vs. REF at the locus (see the Methods section for details).

### 2.2 Data description

We applied eQTLsingle on a published scRNA-seq dataset of glioblastoma multiforme (GBM) samples[22] (referred to as GBM dataset below). GBM is one of the most common brain cancers with high morbidity and mortality. The dataset also includes scRNA-seq data of gliomasphere cells. Scientists often use gliomasphere cell lines as a stem cell (CSC) model of glioblastoma cancer. However, gliomaspheres and glioblastomas have different characteristics[23,24]. We took them as two cell types in our experiment and applied our eQTLsingle method on them separately.

We collected 489 single cells from the dataset after quality control (details in Methods), including 363 glioblastoma cells and 126 gliomasphere cells. We detected 3,507 valid SNVs from the dataset. For the eQTL analysis, we used more stringent criteria and selected 396 and 112 SNVs in glioblastomas and gliomaspheres, respectively. For the possible target genes, we focused on studying possible regulation effects of these mutations on genes that are known to be related to brain tumors. We selected 774 brain-tumor-related genes reported in the literature[22] (Supplementary Table S1) and genes nearby the detected mutations above (up/downstream 1Mb bp of mutations) to study possible eQTLs. After quality control, this gave us a total of 784 and 354 candidate target genes in the experiments on glioblastomas and on gliomasphere, respectively. We use these genes and the detected SNVs to look for both cis- and trans-eQTLs in the two cell types. Details of the selection of candidate SNVs and target genes are described in the Methods.

To validate eQTL findings on single cells, we analyzed GBM patients and LGG (low-grade gliomas, some of LGG develop into GBM) patients in the TCGA data and normal brain tissue samples in the GTEx data (details in Methods).

### 2.3 SNVs called from scRNA-seq data are good representations for single cells

To check whether the SNVs detected from scRNA-seq data are reliable, we performed principal component analysis (PCA) on the 3,507 detected SNVs (Supplementary Figure S2a). We see that these two cell types can be separated clearly in the PC space, and the separation is even better than that based on the gene expression profile (Supplementary Figure S2b). The clear separation of the two cell types by the detected SNVs confirms that SNVs called from scRNA-seq data are reliable and informative, and also highlights the necessity of doing eQTL analysis at single-cell level within each cell types. The observed advantage of single-cell SNV data in distinguishing the two cell types can be benefited from the fact that the mutation profiles do not need normalization across cells, implying that the mutation profile may be a more robust representation for single cells.

### 2.4 Overview of eQTL results on GBM with eQTLsingle

We analyzed associations of the detected SNVs with variations of candidate target gene expression in glioblastoma cells and gliomasphere cells separately, including both local effects (cis-pairs) and distal effects (trans-pair). The results are summarized in Table 1. In the glioblastoma cells, we tested 480 candidate cis-pairs of the 396 SNVs and their 322 nearby genes, and detected 107 significant cis-eQTL pairs (Bonferroni-adjusted p-value < 0.01). Only 19 (17.76%) of these eQTL pairs have been reported in the GTEx database. In the gliomasphere cells, we tested 132 candidate cis-pairs of the 112 SNVs and their 102 nearby genes, and detected 33 significant cis-eQTL pairs (Bonferroni-adjusted p-value < 0.01). Only 4 (12.12%) of them have been reported in the GTEx database. The eQTLs reported in the GTEx database were based on bulk data. Our results show that eQTLsingle applying on single-cell RNA-seq data can reveal many eQTLs at single-cell level that cannot be detected with bulk data. Details of the results are given in Supplementary Table S2.

**Table 1.**
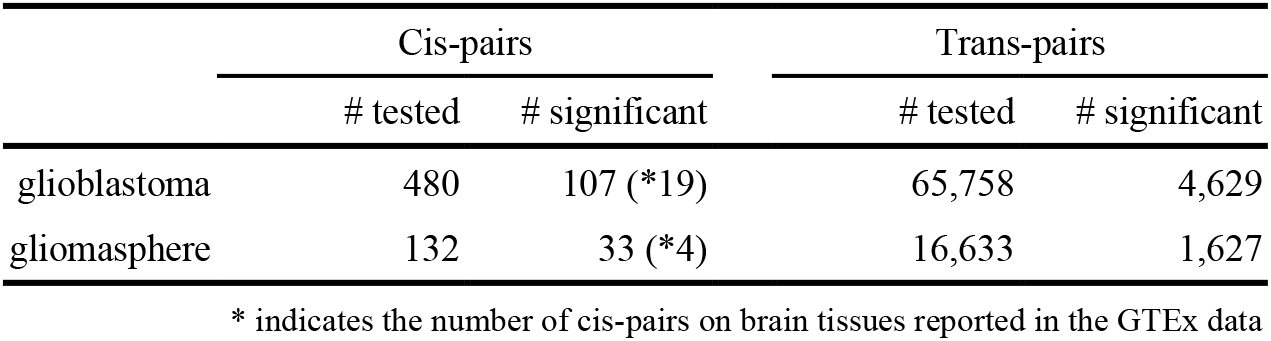
Overview of eQTL analysis result

The GTEx database does not provide systematic results on trans-eQTLs. We applied eQTLsingle on the single-cell data to look for possible trans-eQTLs by taking the pairs of SNVs and selected candidate genes outside the adjacent 1Mb regions of the SNVs as the candidate trans-pairs to be tested. They include 65,758 candidate trans-pairs in glioblastomas and 16,633 candidate trans-pairs in gliomaspheres. Among them, eQTLsingle detected 4,629 pairs of significant trans-eQTLs in glioblastomas, and 1,627 pairs of significant trans-eQTLs in gliomaspheres (Bonferroni-adjusted p-value < 0.01). Details of the results are given in Supplementary Table S2.

The number of new discoveries indicated the importance of analyzing eQTLs in a cell-type-specific manner on single-cell data. We selected some of the discovered single-cell eQTL results as examples to investigate the possible mechanisms behind them.

For cis-eQTL results, we chose eQTL examples of which the variations have been annotated and are in regulatory regions. We focused on variations that have been recorded in dbSNP[25]. In our discoveries, 84 cis-eQTL pairs in glioblastomas and 31 cis-eQTL pairs in gliomaspheres fall in this category. We used HaploReg[26] to get annotations of the variants, including protein binding information from ENCODE[27] and Roadmap Epigenomics[28] projects and effects on regulatory motifs. We narrowed down the choice to variants located on regulatory elements (enhancer, promotor, or loci with histone modifications) or known to influence TF binding, resulting in 51 pairs in glioblastomas and 24 pairs in gliomaspheres. Details of the results are given in Supplementary Table S3. Among them, we chose two example pairs for deeper investigation based on the literature. One is the cis-eQTL pair of SNP rs4758 and target gene PDIA6 discovered in glioblastomas cells. We refer to it as the rs4758-PDIA6 cis-pair for convenience. The other is the cis-eQTL pair of SNP rs3169950 and target gene PSMB6 discovered in gliomasphere cells, referred to as the rs3169950-PSMB6 cis-pair.

For trans-eQTL results, we looked for trans-eQTL pairs of which the variation is non-synonymous and could change the structure and function of corresponding proteins. The numbers of detected trans-eQTLs pairs falling into this category are 377 and 226 in glioblastomas and gliomaspheres, respectively. We analyzed the possible effect of variant on protein function using SIFT[29] and found 78 trans-pairs in glioblastomas and 16 trans-pairs in gliomaspheres with variations that are deleterious for protein functions. Details of the results are given in Supplementary Table S3. We chose the trans-eQTL pair of SNP rs2278161 and target gene C1R in glioblastomas as an example for further investigation, and refer to it as the rs2278161-C1R trans-pair.

### 2.5 rs4758-PDIA6: a cell-type-specific eQTL of different regulatory effects in glioblastoma and gliomasphere cells

The cis-pair rs4758-PDIA6 is a significant eQTL in glioblastomas (Bonferroni-adjusted p-value=1.30 × 10^-4^). The SNP rs4758 is located in the exon region of PDIA6 on chromosome 2p25.1 (Figure 2a), with reference allele cytosine (C) and alternative allele of thymine (T) on the locus. This variation is synonymous in the amino-acid coding. The target gene PDIA6 encodes a member of the protein disulfide isomerase (PDI) family that catalyzes protein folding[30]. In the glioblastoma cells of the GBM dataset, 39.81% are of the REF type and 60.19% are the ALT type. Glioblastoma cells of the ALT type have significantly higher expression level of PDIA6 than those of the REF type (Figure 2b).

**Figure 2.**
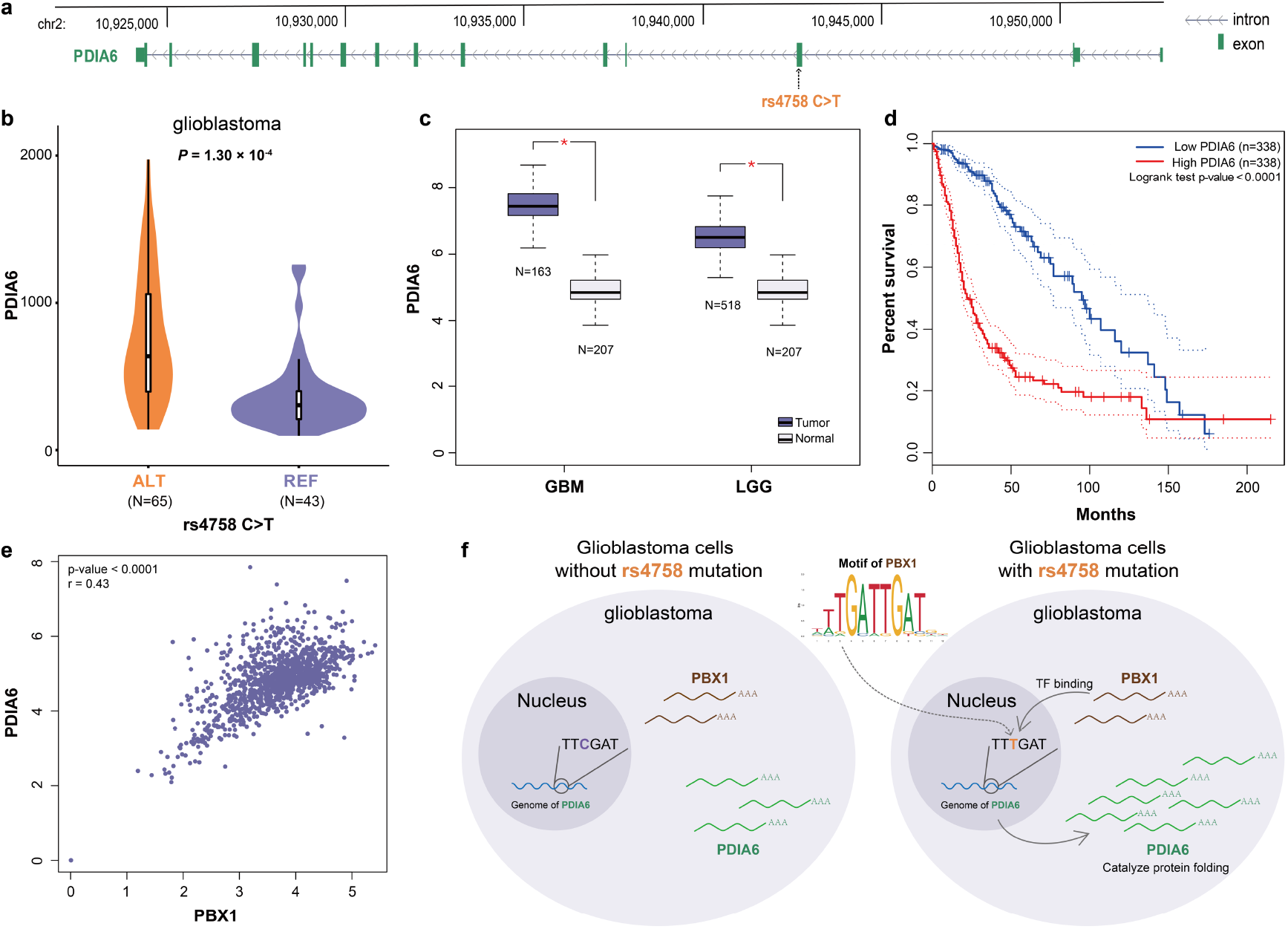
Discoveries on the rs4758-PDIA6 cis-eQTL pair in glioblastomas. **(a)** Genomic location of gene PDIA6 and the SNP rs4758. The top panel is the coordinate of genome. The middle panel depicts the structure of PDIA6. The bottom panel shows the position of variation rs4758. **(b)** PDIA6 expression levels of cells of the ALT and REF genotypes at this locus in glioblastomas (Bonferroni-adjusted p-value=1.30 × 10^-4^). **(c)** Comparison of PDIA6 expression between tumors (GBM and LGG) and normal brain tissues based on the TCGA and GTEx data. Red star indicates the one-way ANOVA test p-value≤ 0.05. **(d)** Kaplan-Meier survival plot of GBM and LGG patients in the TCGA data stratified by gene expression levels of PDIA6. Dotted lines denote 95% confidence interval. **(e)** Gene expression correlation of PBX1 and PDIA6 of brain tissue samples in GTEx. Pearson’s correlation is 0.43 (p-value<0.0001). **(f)** A hypothetical model of the regulation of the cis-eQTL rs4758 on PDIA6 expression. The variation at rs4758 increases the TF-binding affinity of PBX1, which upregulates the transcription of PDIA6. The target gene PDIA6 can catalyze protein folding and enhances tumor growth.

PDIA6 has been known to be essential for tumor growth and inhibition. Knockdown of PDI genes can make cancer cells hard to meet the protein-folding demands during the high-rate synthesis and therefore inhibit cell growth[30–32]. It has been reported that PDIA6 contributes to GBM tumor growth and inhibition of PDIA6 reduces GBM migration and invasion[31,33,34]. We compared the expression of PDIA6 across GBM, LGG, and normal brain tissues in the TCGA and GTEx data, and found that PDIA6 has significantly higher expression levels in GBM and LGG tumors (Figure 2c). Among the GBM and LGG patients, we found that those with higher PDIA6 expression levels have worse overall survival (Figure 2d, see Methods for the detail). These observations show that PDIA6 could enhance GBM progressions.

We found that the variation at rs4758 may increase the affinity of transcription factor PBX1 for PDIA6, and then upregulate PDIA6. Histone markers such as H3K4me1 have been found on this locus in brain tissues[28]. We applied the 15-state ChromHMM[35] model and found that the locus is predicted as an enhancer regulatory element. We analyzed the possible effect of the variation at this SNP on possible TF bindings using HaploReg[26], and found that its variation from the reference type (C) to the alternative type (T) increases affinity of the transcription factor PBX1 binding on this locus according to the position weight matrix (PWM) [36] (Figure 2f). To confirm the relation of PBX1 and PDIA6, we checked their expression in the TCGA and GTEx data. We found that the expressions of PBX1 and PDIA6 have a significant positive correlation of brain tissues from GTEx data (Pearson’s correlation coefficient = 0.43, p-value<0.0001, Figure 2e). These analyses suggested that the mechanism behind the discovered cis-eQTL pair rs4758-PDIA6 is that, the variation at rs4758 upregulates PDIA6 by increasing binding affinity of transcription factor PBX1, as shown in the hypothetic model of Figure 2f.

We checked the association of the SNP rs4758 with the expression of PDIA6 in the gliomasphere cells and found that this pair is not significant at all in this cell type (Bonferroni-adjusted p-value=1, Supplementary Figure S3a). This led to the question why the regulation pathway discovered in glioblastomas is no longer valid in gliomaspheres. We checked the expression levels of the transcription factor PBX1 in the gliomasphere cells and glioblastoma cells, and found that its expression is significantly lower in gliomaspheres (Bonferroni-adjusted p-value=3.47 × 10^-4^, Supplementary Figure S3b). We hypothesized that the low expression of PBX1 may cancel out the impact of variation rs4758 on the binding of this TF and thereby interrupt the regulatory effect on PDIA6 (Figure 2f). The cis-eQTL effect of rs4758 on the target gene PDIA6 has been reported on several tissues like testis and skins in GTEx[37], but no reports on brain tissues. We inferred that different cell types of brain tissues or GBM tumors have different PBX1 expression levels, and therefore this eQTL effect of rs4758 on PDIA6 cannot be found in bulk data. This highlights the importance of doing single cell eQTL analysis separately in different cell types.

### 2.6 rs3169950-PSMB6: a cis-eQTL near the transcription start site that regulates the gene expression in gliomaspheres

The cis-eQTL pair rs3169950-PSMB6 is very significant in gliomasphere cells (Bonferroni-adjusted p-value=2.93 × 10^-12^, Figure 3a). The SNP rs3169950 is located in the first exon of the gene PSMB6 on chromosome 17p13.2, near its transcription start site (shown in bottom of Figure 3e). The reference allele at this locus is guanine (G) (the REF type) and the alternative allele on this locus is adenine (A) (the ALT type). This variation is synonymous in the amino-acid coding. The target gene PSMB6 encodes the protein Proteasome 20S Subunit Beta 6, a part of proteasome 20S, which participates in cell cycle regulation through ATP/ubiquitin-dependent proteolytic degradation of transcription factors[38]. In the gliomasphere cells of GBM dataset, 64.13% are of the REF type and 35.87% are of the ALT type. The gene PSMB6 has significantly higher expression level in gliomasphere cells with the ALT type than those with the REF type (Bonferroni-adjusted p-value=2.93 × 10^-12^, Figure 3a).

**Figure 3.**
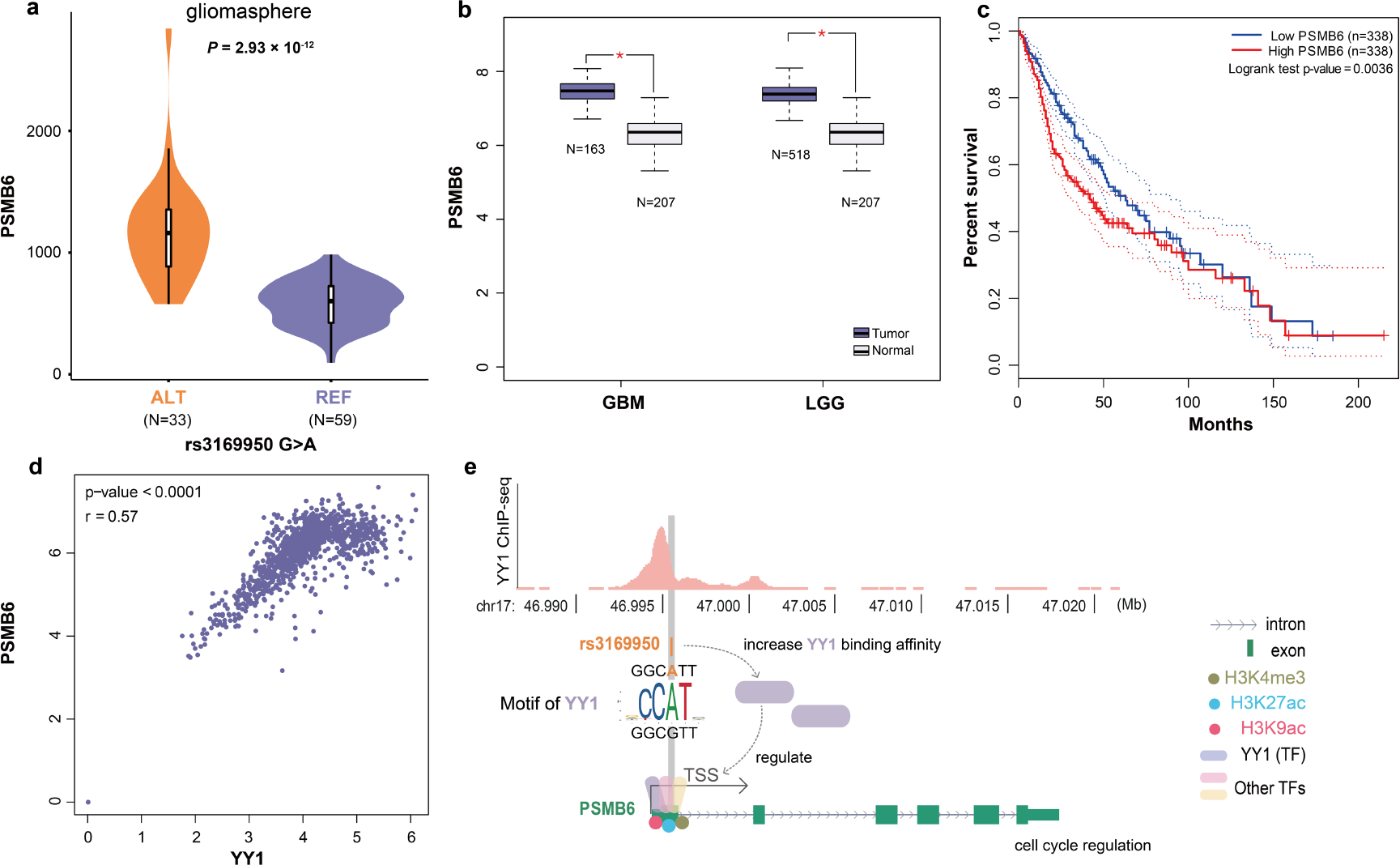
Discoveries on the rs3169950-PSMB6 cis-eQTL pair in gliomaspheres. **(a)** PSMB6 expression levels of cells of the REF/ALT type at rs3169950 in gliomaspheres (Bonferroni-adjusted p-value=2.93 × 10^-12^). **(b)** Comparison of PSMB6 expression between brain tumors (GBM and LGG) and normal brain tissues based on the TCGA and GTEx data. Red star indicates one-way ANOVA test p-value≤ 0.05. **(c)** Kaplan–Meier survival plot of GBM and LGG patients in the TCGA data stratified by gene expression levels of PSMB6. Dotted lines denote 95% confidence interval. **(d)** Gene expression correlation of YY1 and PSMB6 of brain tissue samples in GTEx. Pearson’s correlation coefficient is 0.57 (p-value<0.001). **(e)** ChIP-seq signals of transcription factor YY1 on PSMB6 and a hypothetic model of the regulation of rs3169950 on PSMB6 expression. The upper part shows the scale of peaks from ChIP-seq data of YY1. The genomic coordinates are based on hg19. The middle part shows location of rs3169950 and nearby sequences. The lower part shows histone marks and TFs on PSMB6 and the hypothetical regulatory mechanisms.

PSMB6 and its family members contribute to cell cycle regulation[38] and tumor angiogenesis[39,40]. It has been revealed that PSMB6 was upregulated in hypoxia[41], lung cancer[42], and thyroid cancers[43]. In addition, the gene PSMB8 encoding another part of proteasome 20S subunit, is found involved in tumor angiogenesis and is a candidate biomarker in GBM[39,40]. The clinical benefit of proteasome inhibition treatment is also proven in glioblastoma[44]. To confirm the influence of PSMB6 on GBM, we analyzed expression of PSMB6 across GBM and LGG patients in the TCGA data and normal brain tissues in the GTEx data. We found the expression level of PSMB6 is significantly higher in GBM and LGG tumors than in normal brain tissues (Figure 3b). In addition, we found brain tumor patients with high PSMB6 expression had worse overall survival (Figure 3c). These findings illustrate that PSMB6 is a critical factor in glioma tumorigenesis.

The SNP rs3169950 is located near the transcription start site of PSMB6 and the variation on this locus may influence the transcription process. We applied HaploReg[26] and ChromHMM[35] to analyze this variant. Histone marks including H3K4me3, H3K27ac, and H3K9ac are found near this region[28]. Meanwhile, the 15-state ChromHMM model also predicts it is in the active transcription start site (TssA). This indicates the region is an active regulatory element. We further did database searching and found several TFs (YY1, TAF1, and USF1) act on this region in many regulatory processes (Supplementary Figure S4). The ENCODE ChIP-seq data of YY1 in this region in the SK-N-SH brain cell line is shown in the top panel of Figure 3e. We observed that the expression level of transcription factor YY1 and the target gene PSMB6 have a strong correlation of brain tissues in GTEx data (Pearson’s correlation coefficient=0.57, p-value<0.001, Figure 3d). PWM analysis shows that the variation at rs3169950 could increase YY1 binding affinity (Figure 3e). Thus, we inferred the variation at SNP rs3169950 can influence the binding of TFs such as YY1 on this transcription start region and regulate transcription level of PSMB6 further (Figure 3e).

This cis-pair has low read coverage in the glioblastoma cells in the GMB dataset, and we cannot detect whether it is also significant in those cells.

### 2.7 rs2278161-C1R: a trans-eQTL that alters the protein structure of the target gene in glioblastomas

The trans-eQTL pair rs2278161-C1R is found significant in glioblastomas (Bonferroni-adjusted p-value=3.72 × 10^-3^, Supplementary Figure S5a). The variation is located in the exon area of CNDP2 on chromosome 18q22.3. The reference allele is thymine (T) and the alternative allele is cytosine (C). The target gene C1R is located on chromosome 12p13.31. In the glioblastoma cells of the GBM dataset, 65.71% are of the REF type and 34.29% are of the ALT type. Glioblastoma cells with the ALT type at rs2278161 have significantly higher expression level than those with the REF type.

C1R encodes a subcomponent of complement C1r which is involved in the complement system[45], a part of the immune system. It has been reported that the complement system C1 complex activation is essential for glioblastoma tumorigenesis and glioma stem-like cells maintenance[46,47]. Tumor-cell-derived C1r and C1q have also been reported to promote cell growth in cutaneous squamous cell carcinoma[48,49] and to accelerate glioma cell migration via upregulating expression of several matrix metalloproteinases (MMPs) [47]. We compared expression of C1R across GBM and LGG tumors with normal brain tissues using the TCGA and GTEx data, and found that C1R has a significantly higher expression level in GBM and LGG tumors (Supplementary Figure S5b). We also found GBM and LGG patients with higher C1R expression had worse overall survival (Supplementary Figure S5c). These findings indicated C1R is a risk factor for gliomas.

The variation at rs2278161 in an exon of CNDP2 is non-synonymous, which can cause amino acid sequence change on CNDP2 protein from the large-size aromatic amino acid Tyrosine (Y) to medium size polar amino acid Histidine (H) at position 126 (Supplementary Figure S5f). This locus is conserved reported by Mutfunc[50] and the variant is predicted deleterious by SIFT[29]. We found that this locus is close to metal binding sites such as position 132 and position 167 in 3D space (Supplementary Figure S5f, Uniprot[51] accession code is Q96KP4). Previous research showed that CNDP2 is a metal ion-dependent dipeptidase[52], which indicates the variant near metal binding sites may cause CNDP2 protein dysfunction.

We didn’t found literatures on the direct relationship between CNDP2 and C1R. Previous research have demonstrated that CNDP2 is a functional tumor suppressor via activation of PI3K/AKT Pathway in multiple cancers[53–55]. Xenograft experiments in vivo have shown that CNDP2 could increase the expression of AKT and phosphorylated PI3K[53]. Therefore, dysfunctional CNDP2 protein may inhibit the PI3K/AKT pathway. On the other side, associations between C1R and the PI3K pathway have been revealed in cutaneous squamous cell carcinoma research[47]. Downstream transcription factors of the PI3K pathway such as CREB have also been shown to bind to C1R in multiple cancer cell lines in ENCODE ChIP-seq data[27]. We found that the expression of CNDP2 and C1R are positively correlated in brain tissues from the GTEx data (Pearson’s correlation coefficient=0.36, p-value<0.0001, Supplementary Figure S5d). With the detected eQTL of rs2278161 on CNDP2 with C1R, we speculate that the non-synonymous variation at rs2278161 can cause the dysfunction of the CNDP2 protein, which then influence the expression of C1R.

## 3. Discussion

We developed a new method eQTLsingle to detect single cell eQTLs with scRNA-seq or snRNA-seq data only. Genomic variations are major factors that may influence phenotypes by regulating the expression of target genes. Many of such regulations are specific to cell types or cell states. Conducting eQTL analysis on single-cell data is crucial for revealing the underlying mechanisms, especially for cell types like tumor cells that are rich in genomic variations.

The ideal data for single-cell eQTL analysis are paired sequencing of both the genome and transcriptome of the same single cell. Technologies are being developed toward this direction, but the data obtained with current technology of this type are still of very low coverage and quality, far from sufficient for supporting single-cell eQTL studies. Current single-cell RNA-seq data can be used for eQTL study on the effects of germline variations if samples are collected on multiple individuals. Such “semi” single-cell strategy cannot be used to find the effects of somatic mutations that are especially important for many diseases. eQTLsingle uses the single-cell transcriptomics data for both detecting genomic variations and estimating gene expressions, and thus enables true single-cell eQTL analysis with scRNA-seq or snRNA-seq data only.

Our experiments on the glioblastoma and gliomasphere cells in the GBM scRNA-seq data showed that eQTLsingle can effectively find significant cell-type specific eQTLs that cannot be found by eQTL studied based on bulk data. The example discoveries we dissected not only illustrated the biological significance of the detected single-cell eQTLs, but also demonstrated that the advantage of single-cell-based analysis in resolving the possible regulatory routes and mechanisms underlying the association of variation at the eQTL with the variation in the target gene expression.

We are also aware of the limitation of the proposed method. RNA-seq data only have information of genomic variations in the transcribed regions. Therefore, the possible eQTLs in the noncoding regions and in those lowly expressed genes cannot be detected. Similarly, eQTLsingle works better on scRNA-seq data obtained with full-length sequencing protocols such as SMART-seq. Data obtained with UMI-based technologies only cover a small part of the transcripts. eQTLs that could be detected from such data will be limited. Despite these limitations, the experiments in our study showed that single-cell eQTL using scRNA-seq data only can help to reveal many important regulatory relations and mechanisms that could be missed otherwise, and the proposed eQTLsingle method is a powerful tool for this purpose. We have developed the method as a software package free for academic use and provided it at https://github.com/horsedayday/eQTLsingle.

## 4. Methods

### 4.1 scRNA-seq data pre-processing

The GBM scRNA-seq data is from previous human glioblastomas study[22]. This data was generated with SMART-seq protocol, and was downloaded from the Gene Expression Omnibus (accession code is GSE57872). Sequencing reads (FASTQ files) were aligned to human genome reference (hg19) using STAR 2.7.3a[56] with default parameters. Low-quality cells were filtered as follows: (1) cells with less than 1 × 10^6^ uniquely mapping reads were removed; (2) cells with percentage of uniquely mapped reads less than 60% were removed.

Expression level was calculated using featureCounts 2.0.0[57] with default parameter settings. Gene annotations were downloaded from GENCODE (V19). Cells with less than 3500 genes detected were removed. The data was normalized to CPM (counts per million).

### 4.2 SNV-calling from scRNA-seq data

Mutations were detected from scRNA-seq using GATK[58]. The single-nucleotide variant reference (dbSNP 138) and indel reference (1000G phase1) used here were downloaded from the GATK bundle. Mutations that were detected in at least 40 cells were retained.

Since the loci with no mutations reported may be due to either (1) no mutation or (2) non-sufficient read coverage, we counted the number of reads over each locus in the mutation matrix to determine the condition using bam-readcount[59]. For a locus without mutations reported, (1) genotype type was REF (non-mutated) if the number of reads over it was larger than 5 (denoted as 0 in the mutation matrix); (2) information on genotype type was considered missing data otherwise (denoted as −1 in the mutation matrix). In the following eQTL analyses on some SNV, cells in which the SNV information was missing were filtered out to ensure reliable analyses.

Because of possible unbalanced expression between paternal alleles and maternal alleles of genes, i.e., the allele-specific expression phenomenon in single cells[21], we cannot confidently distinguish homozygous and heterozygous mutation with scRNA-seq data only. Loci with mutations reported along with more than 5 reads covered were considered ALT type (mutated, denoted as 1 in the mutation matrix).

For position *i* in cell *K*, we have genotype *S_ik_*,

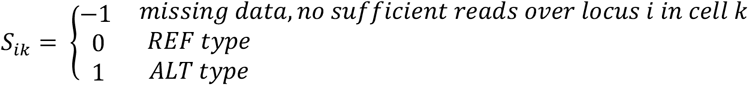

### 4.3 Quality control of candidate target genes

We selected 774 brain-tumor-related genes reported in the literature[22] and genes nearby the detected mutations above (up/downstream 1Mb bp of mutations) to study possible eQTLs. Only genes that were detected in at least 30 cells in both ALT and REF group were kept as candidate target genes for eQTL analysis.

### 4.4 Model scRNA-seq data with the ZINB model

Negative Binomial (NB) model has been widely used for modeling count data[60]. There are excess zeros in scRNA-seq data, and it has been proven zero-inflated negative binomial (ZINB) model can fit scRNA-seq data better[18]. Here we adopted the ZINB model to fit gene expression.

For any r > 0 and p ∈ [0,1], the probability mass function (PMF) of NB distribution is

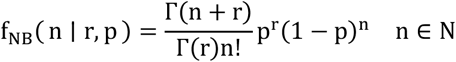

where Γ(∙) is the gamma function.

The ZINB model is a mixture model with parameter θ to depict excessive zeros. For any θ ∈ [0,1], PMF of ZINB distribution for some gene with n read counts in a group of cells is

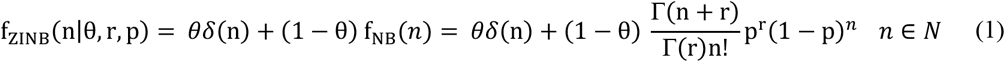

where δ(∙) is the Dirac delta function.

Single cells can be divided into different subsets, such as different cell types or cell states. We divided the GBM dataset into two subsets, i.e., glioblastomas and gliomaspheres. Labels of cells were given by authors of original study[22]. We modeled gene expression within each subset separately.

### 4.5 Discover eQTLs by comparing ZINB parameters between ALT and REF groups

For each detected SNV, we looked for possible eQTL effect on all candidate target genes described above. It was considered candidate cis-pairs if genes were upstream/downstream 1Mb bp of the mutation. Otherwise, it was considered candidate trans-pairs. We discovered eQTLs by comparing gene expression between mutated cell group (ALT) and non-mutated cell group (REF). If some gene expressed differentially in two groups, we considered corresponding SNV is an eQTL affecting the gene (Figure 1).

For discovering the association between genotype of SNV *i* and variations of the expression of gene *j* in a subset, we firstly divided cells belonging to the subset into two groups based on genotype of SNV *i*,

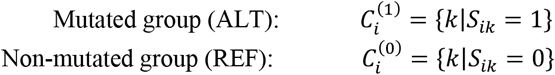

We then modeled gene expression *g_i_* in ALT group 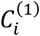 and REF group 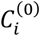 with the ZINB model (equation (1)) separately. ZINB parameter of ALT group 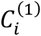 is denoted as 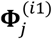 and ZINB parameter of REF group 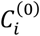 is denoted as 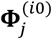:

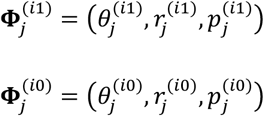

We used 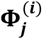 to denote two parameters above together:

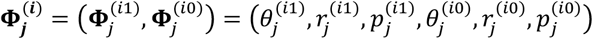

We considered gene *j* expressed differentially in ALT and REF group if any of three elements in 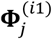 and 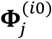 were significantly different. Therefore SNV i is an eQTL affecting gene *j*. We compared differences of ZINB parameters by testing null hypothesis:

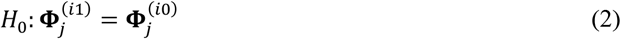

Hypothesis testing worked as follows:
Firstly, we estimated 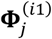 and 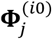 in ALT group 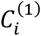 and REF group 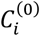 using maximum likelihood estimation (MLE) algorithm without constraints, respectively:

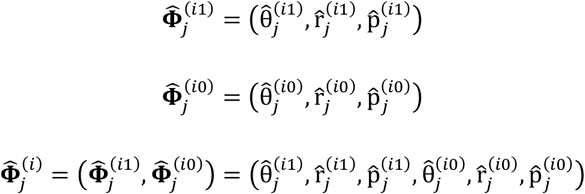

Secondly, we re-estimated 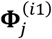 and 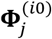 in two groups using MLE algorithm under the constraint (equation (2)). This equals to estimate parameters on both groups 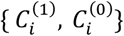, together. Then we got:

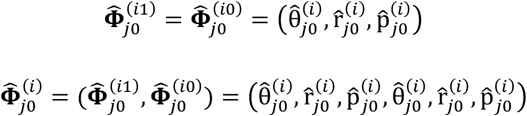

Thirdly, since the models were nested, we applied likelihood ratio test (LRT) to test the hypothesis (equation (2)). The likelihood ratio statistic was:

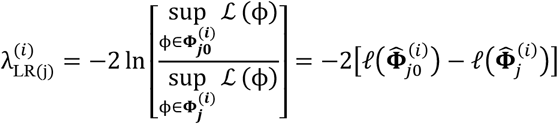

According to Wilks’ theorem[61], under the case the null hypothesis is true, the likelihood ratio statistic 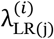 follows chi-square distribution 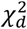,

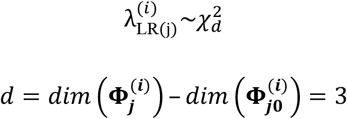

Where *d* is the degree of freedom for 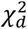.

Therefore, we calculated p-value:

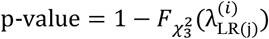

Where 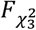 is cumulative distribution function of 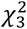

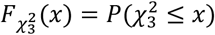

We used Bonferroni correction to correct for multiple testing on candidate cis-pairs and trans-pairs in each subset separately. The candidate pairs were considered significant if Bonferroni-adjusted p-value < 0.01.

### 4.6 TCGA and GTEx data analysis

We applied GEPIA[62] to study the gene expression data from TCGA and GTEx projects, including comparing gene expression between tumors and normal tissues, survival analysis of tumor patients, and correlation analysis between expression levels of genes.

For comparing gene expression between tumors and normal tissues, we set GBM patients (n=163) and LGG patients (n=518) in TCGA as tumor group and appropriate normal brain tissue samples (n=207) in GTEx as normal group. Appropriate normal brain tissue samples for comparing GBM and LGG tumors were labeled by medical experts from the GEPIA[62] team. We transformed expression data with log2(TPM + 1) for visualization.

For survival analysis, we chose GBM and LGG patients with survival information (n=676) in TCGA. We divided these patients into low-expression group and high-expression group using median value as group cutoff. We adopted logrank test to compare overall survival distributions of the two groups.

For correlation analysis of gene expression levels, we applied Pearson method to calculate correlation coefficient on brain tissue samples (n= 945) in GTEx.

### 4.7 Functional annotation and analysis of variants

We applied HaploReg[26] (v4.1) to explore annotations of variants, including functional consequences, protein binding results, chromatin state segmentations, and effects on regulatory motifs. Functional consequences are obtained from dbSNP database. Chromatin modification data is from ChIP-seq experiments in Roadmap Epigenomics[28] and Encode[27] projects. Chromatin states are predicted with ChromHMM[35] model. This model integrates ChIP-seq data of different histone marks from Roadmap Epigenomics Project and classifies genome into promoter, enhancer, and other regulatory states. Effects on regulatory motifs are analyzed based on position weight matrices (PWMs). PWMs are curated from JASPAR[63], TRANSFAC[64], protein-binding microarray experiments[65] and ChIP-seq experiments[36].

The potential functional consequences of non-synonymous variations on their located genes are predicted by SIFT[29]. We used ANNOVAR[66] (2019Oct24) as a port to get SIFT score for each non-synonymous variation, and it’s considered deleterious if SIFT score is lower than 0.05. We studied function of proteins with UniProt[67] and analyzed sequence conservation on proteins using Mutfunc[50].

## Abbreviations

eQTL: expression quantitative trait locus
scRNA-seq: single-cell RNA sequencing
ZINB: zero-inflated negative binomial
SNV: single nucleotide variant
LRT: likelihood ratio test
GBM: glioblastoma multiforme
CSC: cancer stem cells
PCA: principal component analysis
LGG: low-grade gliomas
TF: transcription factor
ChIP-seq: chromatin immunoprecipitation sequencing
PWM: position weight matrix
TSS: transcription start site
PMF: probability mass function
NB: negative binomial
MLE: maximum likelihood estimation

## Software availability

An open-source software implementation of our eQTLsingle method is available for free academic use at https://github.com/horsedayday/eQTLsingle.

## Authors’ contributions

XZ and TM conceived and designed the study. TM conducted experiments and developed the software. HL and TM carried out the data analysis. XZ, TM and HL wrote the manuscript.

## Acknowledgments

The authors would like to thank Dr. Kui Hua for helpful discussions.

